# Molecular Dynamics Investigation of the Role of Residues D137 and S315 to INH Binding in KatG

**DOI:** 10.1101/245407

**Authors:** Matt Bawn, Rick Magliozzo

## Abstract

Isoniazid (INH) is a front-line anti tuberculosis drug that is activated *in vivo* by the *Mycobacterium tuberculosis (Mtb)* heme containing catalase-peroxidase (KatG) protein. A single mutation in this protein, S315T, is believed to be associated with up to 50% of drug-resistant cases of Mtb and another mutation D137S has recently been shown to exhibit increased INH activation. The present work uses molecular dynamics simulations and molecular modeling approaches to examine INH binding in the wild-type protein as well as the S315T and D137S mutants. The results suggest that INH binding to the heme site in the S315T mutant is restricted due to reduced access to the binding pocket. Conversely in the D137S mutant INH access to the binding pocket is enhanced especially with-regards-to the heme propionate groups. It is proposed that the increased activation of INH in the D137S mutant may be due to the involvement of an electron transfer pathway facilitated by the heme propionate groups.

## Introduction

Then *Mycobacterium tuberculosis* (Mtb) catalase-peroxidase (KatG) protein is a bi-functional enzyme that exists as a homodimer of 80-kDa subunits^1^ (Figure 1). Each subunit binds one heme cofactor, exhibiting a binding pocket environment and sequence homology that classifies the enzyme as a class I peroxidase. KatG also contains a covalently-linked three amino acid adduct in the distal heme pocket, comprised of W107, Y229 and M255 that is not found in monofunctional peroxidases. Mutagenesis studies have indicated that the adduct is necessary for catalase but not peroxidase activity. Existing crystal structures of KatG show adduct formation in each case and a high degree tertiary structure conformity prevailing throughout.

**Figure 1.**
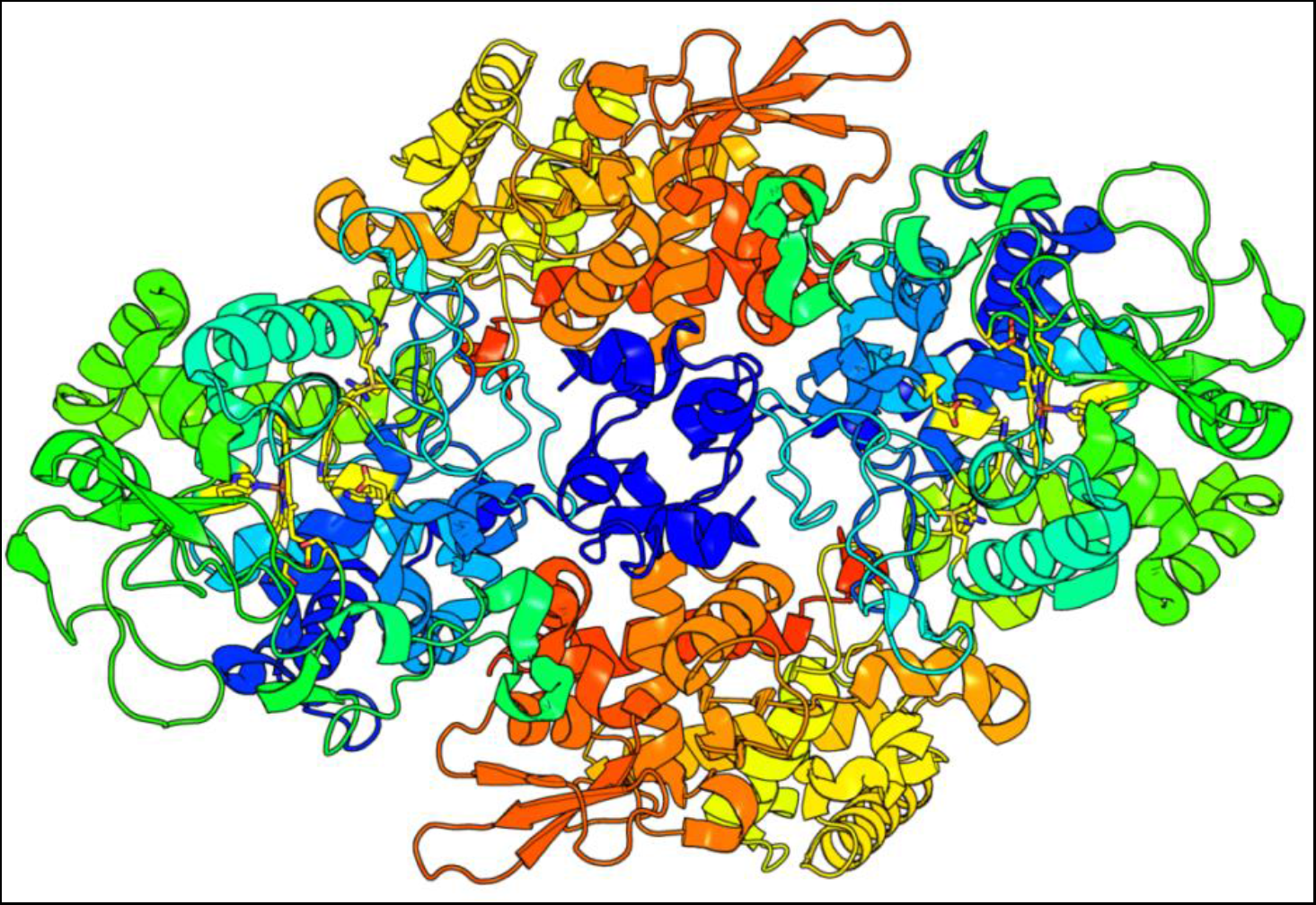
Structure of MTb KatG (from PDB file 2CCA) with binding-pocket amino acid residues and heme cofactors shown as sticks in yellow.

Mtb KatG has also received interest due to its role in the activation of the anti tuberculosis pro-drug isoniazid (INH), particularly with regards to the prevalent S315T mutant in INH inactivation. Isoniazid (INH) is a pro-drug used to combat Mtb worldwide. It is thought that INH is oxidised *in-vivo* by Mtb KatG to form an acetylating species capable of forming an adduct (IN-NAD) upon reaction with NAD+. This adduct is then capable of inhibiting the Mtb enoyl-acyl carrier protein reductase (InhA), disrupting cell wall biosynthesis^2^. Previous MD studies of the S315T mutant of KatG and its role in INH activation have revealed overall increases in the root mean square deviations (RMSD) of the protein backbone in the mutant compared to the wild-type protein but decreased fluctuations for INH binding residues. The authors in each case suggest the increased rigidity of the binding residues hampers INH activity. Our group has recently reported that the D137S mutant of KatG displays a greatly increased catalysis for INH activation^3^. In an attempt to explain this finding the following work utilises molecular modelling to elucidate the role of the D137S mutant of KatG in INH activation.

## Materials and Methods

*KatG Crystal Structures:* The crystal structure of wild-type and S315T KatG used in this study was previously determined by our group (PDB ID 2CCA and 2CCD respectively). The structure of D137S has been recently described^3^. Crystallographic water molecules were removed from structure prior to molecular dynamics simulations. Crystal structures and model structures were manipulated using PyMol (DeLano Scientific LLC).

*Molecular Dynamics:* The molecular dynamics simulations were performed using the software program Gromacs^4^ version 4.5.5 running on the CUNY high performance computing cluster in parallel using 16 nodes. The GROMOS96 43a1 force field was used and heme cofactor was also incorporated into the simulation. The protein was solvated using the SPC/E water model^5^ centered in a dodecahedron with a minimum distance of 1.0 nm from the protein to the box edge (volume 1724 nm^3^). The number of water residues included in the simulations were 51072 (WT) and 50300 (S315T) 51672 (D137S). The net charge of each system was neutralized by the addition of 44 sodium ions to the solvent. Initial structures were relaxed through a 1000 step deepest descent energy minimization to a maximum force of 1000 kJ/mol/nm. The simulation system was then equilibrated at 300 K as a constant number of particles, pressure, volume and temperature ensemble for 100 ps. Equilibrated models were then subjected to MD simulations for 20000 ps using 0.002 ps steps at 300 K. Statistical data generated from the MD simulations is displayed using MatLab (MathWorks MA).

*Ligand Binding*: Cut-down structures from MD simulations at 17 ns were used without the presence of water molecules. All residue atoms within 1.5 nm of the heme were selected and had hydrogen’s added. The structure of INH was obtained from the crystal structure 3N3S^6^. Hydrogen’s were added and the structure was energy optimized using ArgusLab^7^. The same program was then used to determine the INH binding sites to the heme group for a 6 × 6 × 6 nm cube with a 0.4 grid resolution centered on the heme group for each of the cut-down structures. The ArgusDock engine was used with the INH molecule set as a flexible ligand and calculation comprised 150 poses.

*Binding Pocket Volumes:* Heme binding pocket volumes and environments were determined using the CASTp (Computed Atlas of Surface Topography of proteins) server^8^. This server provides an online resource for locating and measuring potential binding-pocket regions within the three-dimensional structure of proteins. A probe radius of 0.14 nm was selected and the heme group was included.

## Results

*Molecular Dynamics:* The trend in RMSD and radius of gyration of the protein backbone of each MD simulation are shown in Figure 2. The radius of gyration indicates the degree of compactness and shows that each structure becomes more compact throughout the course of the 20 ns simulation. The Root Mean Squared Fluctuation (RMSF) of each residue in the structures during the MD simulation is also shown Figure 3. The residues with an RMSF greater than 1.0 nm are indicated on the figure. The distance distribution between residues 137 and 315 that comprise the “gate” to the binding pocket over the complete course of the MD simulation is shown in Figure 4. The mean distance for each structure is 0.803, 0.819 and 0.836 nm for S315, WT and D137S respectively. It should be remembered that H-atoms are not included in the simulation.

**Figure 2.**
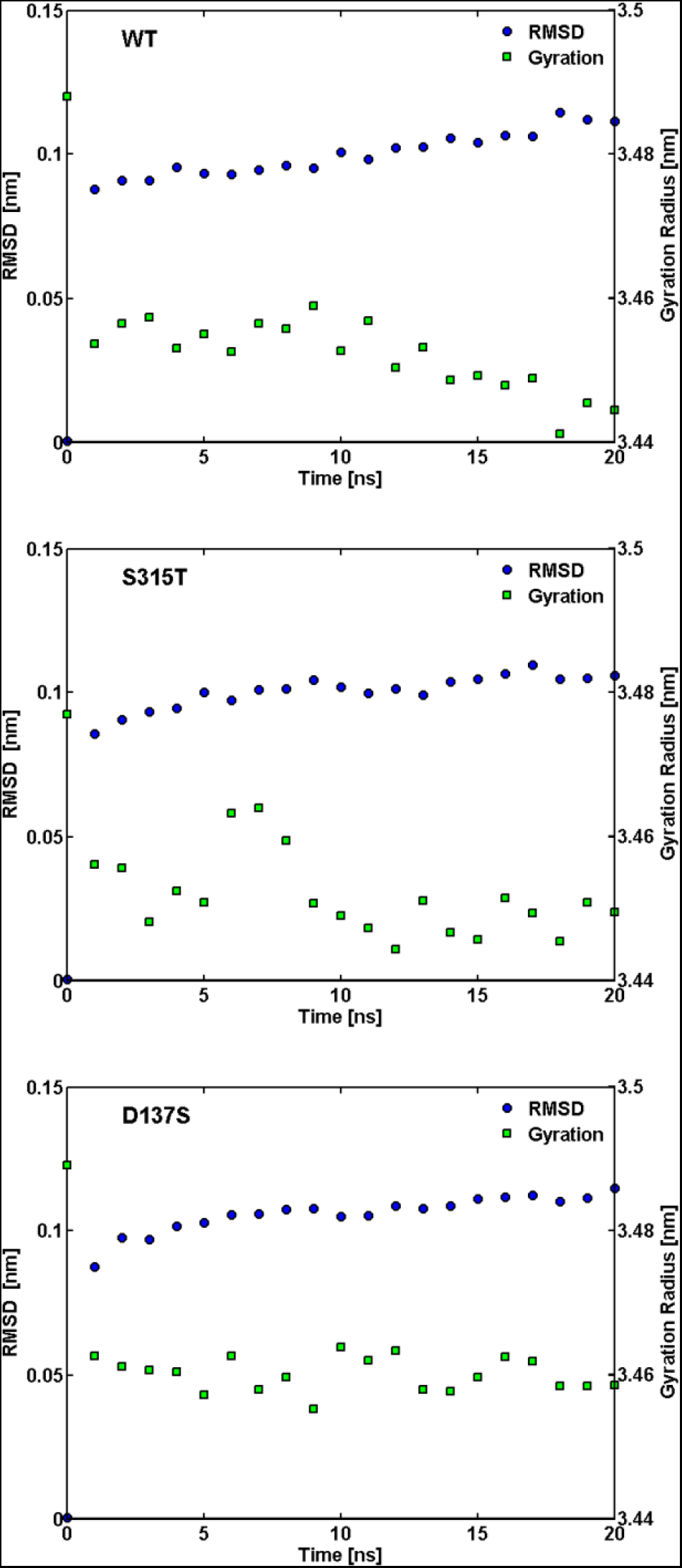
RMSD of protein backbone of MD simulation compared to initial structure plotted against radius of gyration for WT (top), S315T (middle) and D137S (bottom).

**Figure 3.**
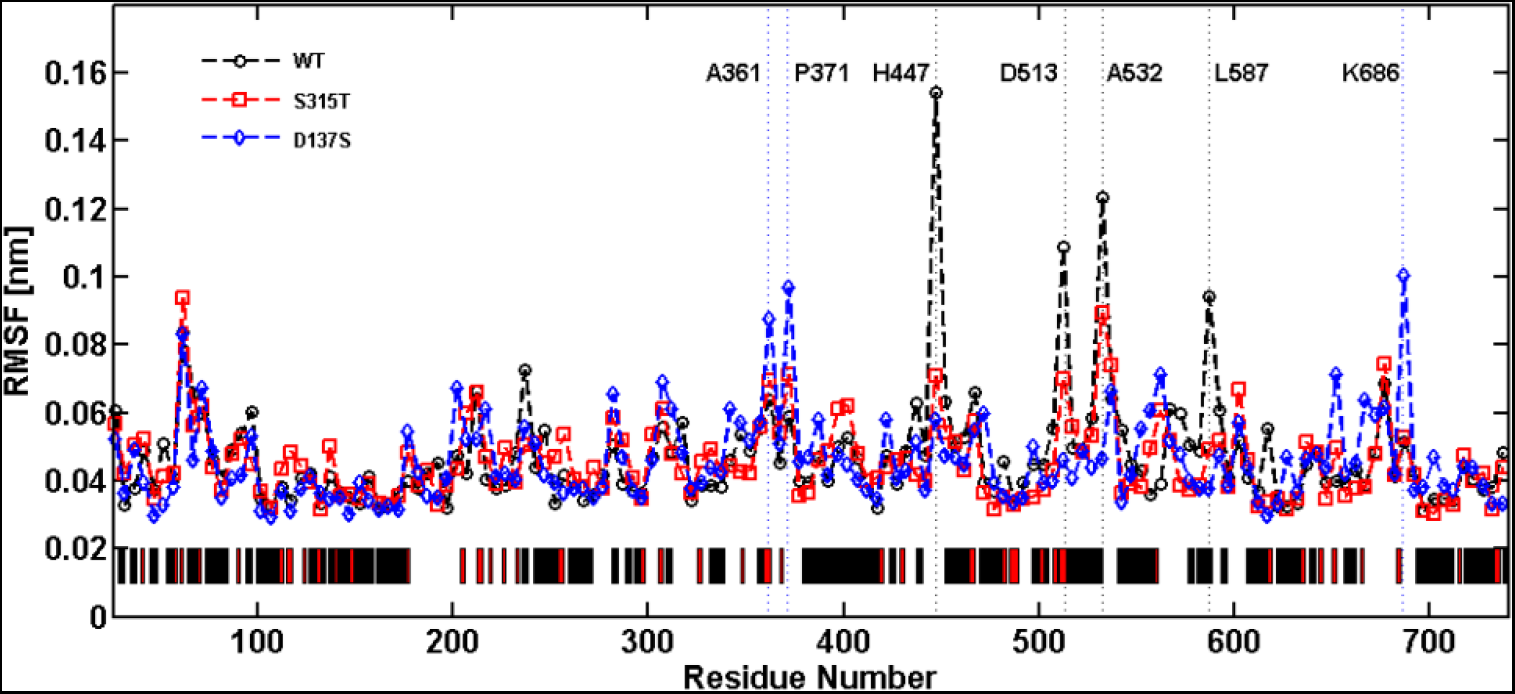
RMSF during 20 ns MD run of WT, S315T and D137S KatG. The bar at the bottom of the figure displays α-helix and loop-region secondary structure (black and red respectively)

**Figure 4.**
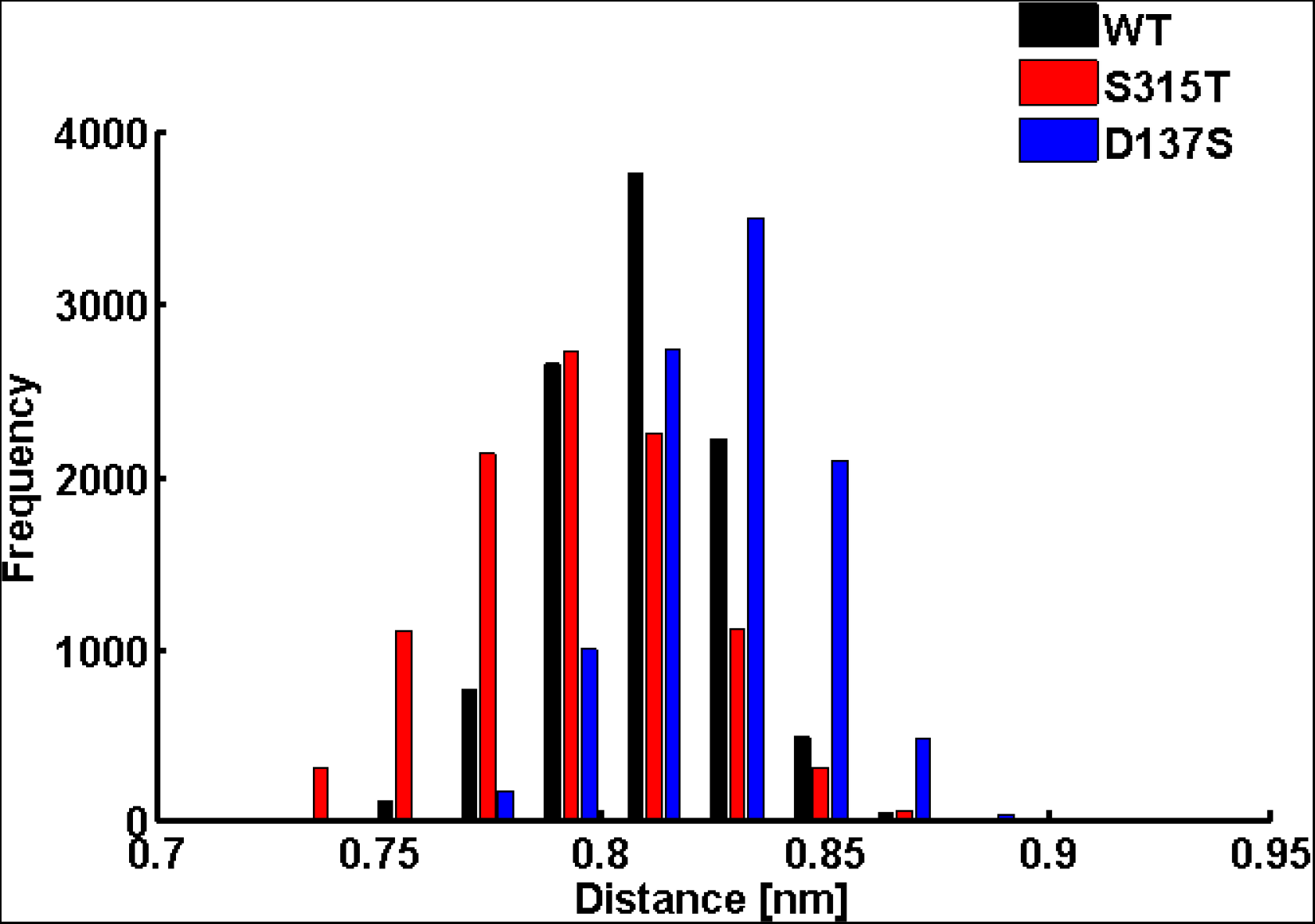
Distance between residues 137 and 315 over the course of the MD simulation for WT S315T and D137S, plotted as histograms of frequency distributions.

*Ligand Binding:* The calculated locations of INH binding to cut-down structures of KatG are shown in Figure 5. It is clear that there is a marked difference in the calculated binding location of the INH molecule in each model. The calculated binding energies for each case are: −7.84, −7.60 and −6.80 kcal/mol for the WT, D137S and S315T models respectively. For the WT INH is shown bound near the heme δ-edge, whilst for the D137S mutant the INH molecule is seen near the heme α-edge. Interestingly in the S315T mutant INH is indicated to be outside of the heme binding pocket.

**Figure 5.**
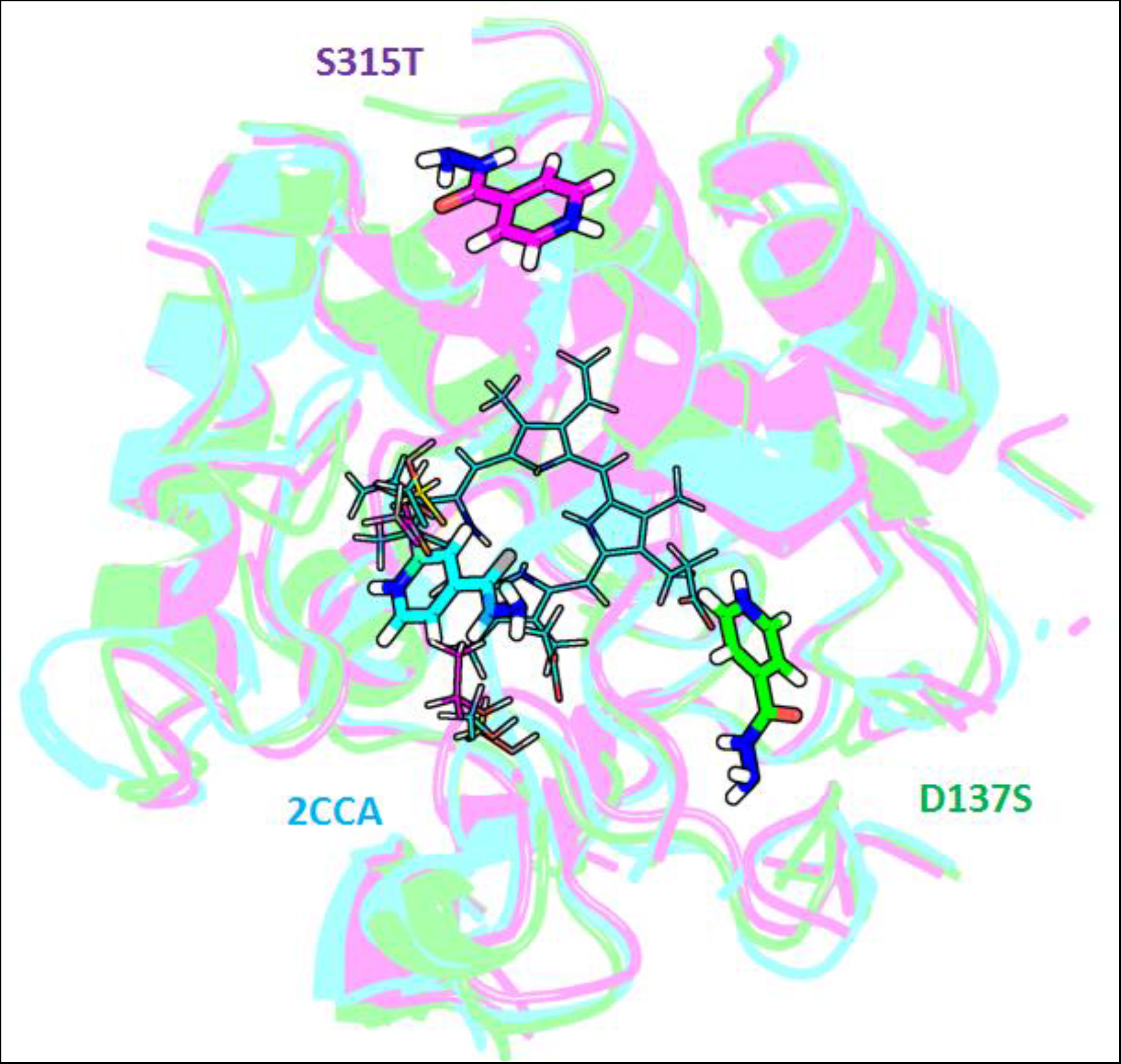
INH docking into the heme binding pocket of 2CCA, 2CCD and D137S structures after 17 ns. Porphyrin propionates pointing towards bottom of picture, residues 137 and 315 on left of heme. To simplify the figure only the heme from the WT (2CCA) is shown.

*CAST_p_ Binding Pocket Volume:* Wire mesh profiles of the determined binding pockets for each crystal structure are shown in Figure 6. The general trend in terms of volume of binding pocket is D137S > WT > S315T. For D137S and WT structures the accessible binding pocket contains the complete heme group, however, for S315T the binding pocket contains just the access gate region (the area between residues 137 and 315) and a small area of the heme δ-edge.

**Figure 6.**
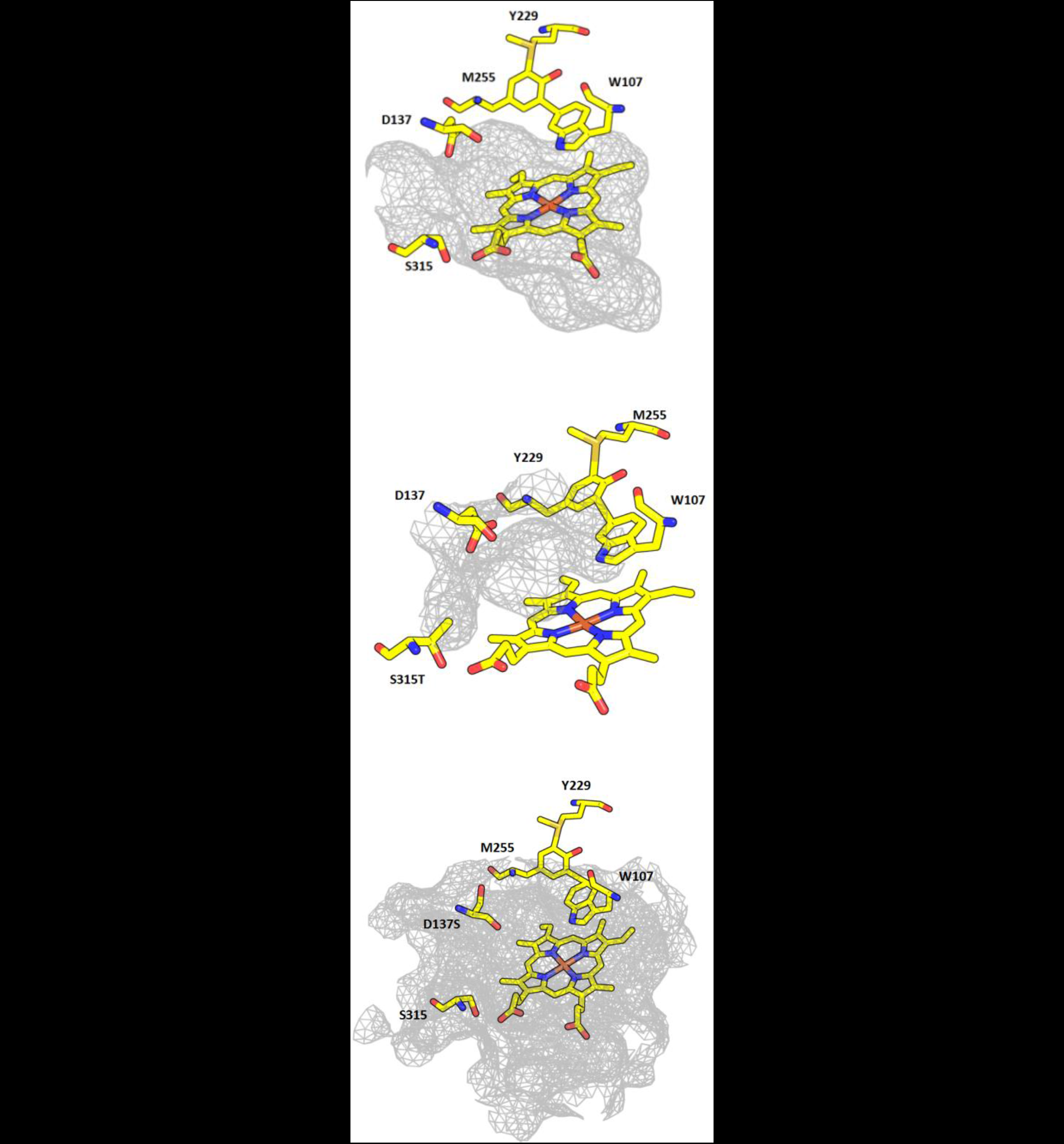
Binding pocket volumes calculated by CASTp of WT, S315T and D137S (top to bottom)KatG structures.

## Discussion

*INH Binding to KatG:* The distance between the gate residues 137 and 315 during the course of the MD simulations revealed in Figure 4 clearly support the assertion made in our recent paper^3^ that INH activation may be hindered in the S315T mutant by restricted access to the binding pocket by showing a reduced gate diameter between D137 and T315 in the S315T mutant. The larger gate distance seen in the D137S mutant is in agreement with the experimental data for enzymatic activation of INH previously reported. The CASTp determination of the extent of the binding pocket Figure 6 also corroborates a restricted access for the INH molecule to the heme in the binding pocket, with the heme in the S315T mutant model being almost completely inaccessible. The ligand binding results reinforce these observations showing the INH molecule to bind near the heme δ-edge in the WT, the heme γ-edge in the D137S and outside of the heme binding pocket in the S315T models. INH binding has also previously been shown mostly at the heme α but also at the γ edges in Ascorbate (APX) and Cytochrome *c* peroxidase (CcP)^9^. Comparisons of the binding locations in APX and CcP to those found in this work are shown in Figure 7, it is clear that there is a strong agreement to those of the WT and D137S models. Metcalfe *et al* found a water mediated hydrogen bond network linking INH to the propionate group of the heme for the γ-edge bound molecule in APX.

**Figure 7.**
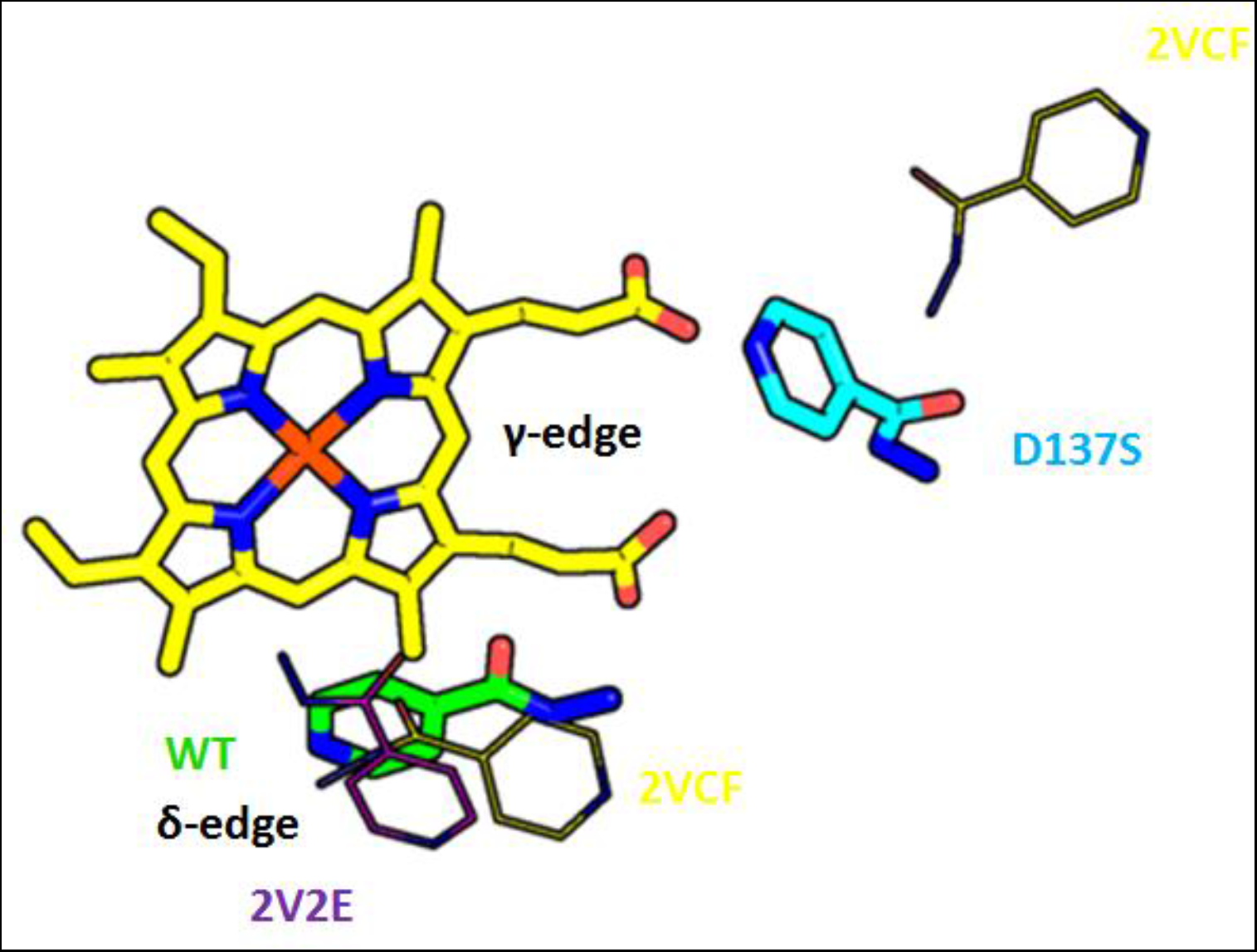
Calculated INH Binding sites of WT and D137S MD models. Compared to binding sites from INH-bound crystal structures of Ascorbate peroxidase and Cytochrome c peroxidase (2VCF and 2V2E). The heme δ and γ-edges are also indicated.

Several experimental and theoretical studies have identified an active role of the heme propionate groups in electron transfers. Guallar *et al* have performed calculations on the role of these groups in peroxidises that indicated that the carboxylic group could play a role in electron transfer processes the mechanism of which they termed the propionate e-pathway^10^. This may therefore suggest that the enhanced enzyme activity associated with INH activation in D137S is facilitated by the involvement of the heme propionate group in a similar e-pathway process in turn aided by an increased binding pocket access and volume when compared to the WT protein.

## Acknowledgements

This research was supported, in part, by a grant of computer time from the City University of New York High Performance Computing Center under NSF Grants CNS-0855217 and CNS-0958379.

